# Spinal cord injury: a systematic review and meta-analysis of microRNA alterations

**DOI:** 10.1101/2023.08.05.551159

**Authors:** Seth Stravers Tigchelaar, Harsh Wadhwa, Maya B. Mathur, Zihuai He, Suzanne Tharin

**Affiliations:** Department of Neurosurgery, Stanford University Medical Center, Stanford, CA, USA; Department of Pediatrics, Stanford University Medical Center, Stanford, CA, USA; Department of Neurology, Stanford University Medical Center, Stanford, CA, USA; Division of Neurosurgery, Palo Alto VA, CA, USA

## Abstract

Spinal cord injury (SCI) often results in life-long disability, lost wages, reduced quality of life, and high economic burden. A bottle neck in the performance of clinical trials exists, in part, due to a lack of diagnostic and prognostic markers of injury severity and neurologic recovery. In addition, while many interventions show promise in preclinical animal models, there are currently no neurorestorative treatments for SCI patients. The development of objective biological markers (biomarkers) and molecular targets for novel treatments for SCI represent two urgent unmet clinical needs. MicroRNAs (miRNAs) represent promising molecules as objective and informative molecules of injury severity and recovery and as potential therapeutic targets. miRNAs are small, stable, regulatory RNA molecules that are evolutionarily conserved across species. miRNAs represent powerful predictors of pathology, particularly with respect to neurologic disorders. Here, we present a systematic review and meta-analysis investigating the conserved inter- and intra-species miRNA changes that occur post-SCI and provide a comprehensive resource for the SCI community. Our analysis identifies a robust set of miRNAs that are involved in the pathophysiologic processes activated in response to SCI.

## Background

In the last four decades, improvements in medical, surgical, and rehabilitative care have increased the quality of life and extended the life expectancy of individuals with SCI – once considered an imminently fatal condition [45]. However, while management has improved, there are no therapies for SCI that have demonstrated convincing neurologic benefit in large-scale clinical trials [14, 42–44, 46]. There are, therefore, urgent unmet needs for the preclinical scientific development of novel therapeutic strategies and for the subsequent clinical validation of these treatments in human trials.

miRNAs hold great promise to underlie novel treatment strategies for SCI. miRNAs are small (22 nucleotide), noncoding RNAs that regulate at least 30% of all protein coding genes [28]. miRNAs are highly expressed in the central nervous system (CNS) [33, 35, 39, 41], with several showing CNS-specific expression [38]: nearly 80% of detected miRNAs are expressed in the adult rat spinal cord [29]. Microarray analysis, real-time PCR, and *in-situ* hybridization have identified 44 miRNAs more than threefold enriched in the brain or spinal cord [33]. Some miRNAs are expressed in a cell type-specific manner in the CNS, with specific miRNAs being expressed in neurons [37, 38], astrocytes [25, 36], and oligodendrocytes [31]. We have shown that miRNAs are required for the development and possibly the evolution of the corticospinal system [6], injury to which results in paralysis in SCI [9].

We and others have shown that miRNA levels in cerebrospinal fluid (CSF) and blood serum are specifically altered in acute SCI in a severity-dependent fashion, both in humans [11], and in animal models [15, 29]. While less is known regarding miRNAs in chronic SCI, miRNAs are differentially altered in chronic SCI patients undergoing active exercise regimes, compared to sedentary patients [10]. miRNAs are promising biomarkers of injury severity, and, by implication, recovery and response to treatment [7, 8, 22] due to their regional abundance, specific developmental requirements, and altered levels following SCI. miRNAs represent promising therapeutic targets for CNS injury and disease, given their intimate involvement in the development and injury of the relevant circuitry, their ability to cross barriers and membranes, and their potential for rapid transition from bench to bedside [18, 23].

To inform the next stage of miRNA biomarker and preclinical therapeutic interventions, this study provides a comprehensive and systematic understanding of the disparate existing datasets on miRNA changes in preclinical models of SCI as well as human patients with traumatic SCI. There has been no meta-analysis of miRNA changes in the setting of SCI that includes the results of recent human trials – a critical gap in the integrated understanding of the pathophysiological response of miRNAs to SCI. The present study is a systematic review and meta-analysis of the current literature, in which we provide a comprehensive analysis of the miRNA response to SCI. We establish the complete set of miRNAs implicated in the pathophysiology of SCI and integrate this experimentally validated miRNA dataset to identify temporal and tissue-targeted patterns of expression, setting the stage for significant advancement in SCI research at the bench. We identify a group of evolutionarily conserved, SCI-relevant miRNAs, high-value targets for potential future biomarkers and therapies.

## Methods/Design

### Protocol registration and standard reporting

For the preparation and development of this study, we followed the checklist provided by the Preferred Reporting Items for Systematic Reviews and Meta-Analyses Protocols (PRISMA) [3]. The protocol for the systematic review and meta-analysis has been registered through PROSPERO: CRD42021222552, and peer-reviewed and published for timely dissemination [1].

### Information sources and search strategy

Studies investigating miRNA alterations in all species of animal models of SCI and human studies were identified from PubMed, Embase, and Scopus. Our search strategy was developed in consultation with an expert librarian and information specialist in systematic reviews.

Our PubMed search strategy was as follows: "MicroRNAs"[Mesh] or "RNA, Untranslated"[Mesh] OR "untranslated rna"/exp OR "microrna*"[tw] OR "mirna*"[tw] OR "micro rna*"[tw] OR "mir"[tw] OR "non coding rna*"[tw] OR ncrna*[tw] OR "non protein coding rna*"[tw] OR "noncoding rna*"[tw] OR "untranslated rna*"[tw] or "npcrna*"[tw] or "non translated rna*"[tw] or "non peptide coding rna*"[tw] AND "Spinal Cord Injuries"[Mesh] or "Spinal Cord Trauma*"[tw] or "Traumatic Myelopath*"[tw] or "Spinal Cord Injur*"[tw] or "Spinal Cord Transection*"[tw] or "Spinal Cord Laceration*"[tw] or "Post-Traumatic Myelopath*"[tw] or "Spinal Cord Contusion*"[tw].

Our Embase search strategy was as follows: ’microrna’/exp OR ’untranslated rna’/exp OR ’microrna*’:ti,ab,kw OR ’mirna*’:ti,ab,kw OR ’micro rna*’:ti,ab,kw OR ’mir’:ti,ab,kw OR ’non- coding rna*’:ti,ab,kw OR ncrna*:ti,ab,kw OR ’non protein coding rna*’:ti,ab,kw OR ’noncoding rna*’:ti,ab,kw OR ’untranslated rna*’:ti,ab,kw OR ’npcrna*’:ti,ab,kw OR ’non translated rna*’:ti,ab,kw OR ’non peptide coding rna*’:ti,ab,kw AND ’spinal cord injuries’/exp OR ‘spinal cord trauma*’:ti,ab,kw or ’traumatic myelopath*’:ti,ab,kw OR ’spinal cord injur*’:ti,ab,kw OR ’spinal cord transection*’:ti,ab,kw OR ’spinal cord laceration*’:ti,ab,kw OR ’post-traumatic myelopath*’:ti,ab,kw OR ’spinal cord contusion*’:ti,ab,kw.

Our Scopus search strategy was as follows: ( TITLE-ABS-KEY ( "spinal cord trauma*" OR "traumatic myelopath*" OR "spinal cord injur*" OR "spinal cord transection*" OR "spinal cord laceration*" OR "post-traumatic myelopath*" OR "spinal cord contusion*" ) ) AND ( TITLE-ABS-KEY ( "microrna*" OR "mirna*" OR "micro rna*" OR "mir" OR "non-coding rna*" OR ncrna* OR "non protein coding rna*" OR "noncoding rna*" OR "untranslated rna*" OR "npcrna*" OR "non translated rna*" OR "non peptide coding rna*" ) ).

### Eligibility criteria

The inclusion criteria for this study were as follows: 1. Studies published anytime, 2. Including all species, and sexes with acute, traumatic SCI, 3. Relating to the alteration of miRNAs following SCI, using molecular-based detection platforms including qRT-PCR, microarray, and RNA-Sequencing, 4. Including statistically significant miRNA alterations in tissues, such as spinal cord, serum/plasma, and/or CSF, and 5. Studies with a SHAM surgery group. The exclusion criteria for this study were as follows: 1. Studies not relating to acute, traumatic SCI, 2. Non-English articles, 3. *In vitro*, *ex vivo*, in silico studies, 4. Non-peer-reviewed articles and conference abstracts, 5. Studies focused solely on differential expression of miRNAs in chronic SCI, defined here as greater than seven days post SCI, and 6. Case reports or case series.

### Study selection

All original studies were eligible for inclusion. Two reviewers screened the imported studies using Covidence [47], reading title and abstract, and excluded studies that did not meet the inclusion criteria. At the selection phase, two reviewers independently included studies that met all eligibility criteria. Any discrepancies were resolved by consensus. A PRISMA flow diagram representing the results of study selection are shown in Figure 2.

### Inclusion and exclusion criteria

#### Animals/population

All species and sexes of animal models and studies including human patients were included in this analysis. Any *in vitro* studies, *ex vivo* studies, in silico studies, and polytrauma studies that did not include *in vivo* analysis were not included.

#### Intervention/exposures

All studies including animal models and human studies of SCI were included. Any study of CNS injury not including the spinal cord was not included.

#### Comparator group

All studies with the inclusion of a comparator or control group (group without SCI) were included. Any study without a control group was not included.

#### Outcome measures

All studies including the detection of miRNAs and their alteration, in spinal cord parenchyma, blood, plasma/serum, or CSF, in response to SCI and within seven days were included. Any study that did not report the direction of change of miRNA in response to SCI was not included.

### Data extraction and data items

Data were extracted by two reviewers. Data parameters collected included: 1. Author/year, 2. Study title, 3. Study model, 4. Study subjects (species, sex, sample size), 5. SCI model (contusion, transection, ischemic-reperfusion), 6. miRNA detection platform (RT-PCR, microarray, sequencing), 7. miRNA direction and/or magnitude of change, and 8. Reported p-value and/or q-value of each miRNA change. When these values were not explicitly provided, the extraction of statistical data from graphs was performed using the graphical data extraction application, WebPlotDigitizer (Version 4.4) [16].

#### Subject characteristics

Information including species, sex, and age was collected. If numerical values were presented as an interval, then the mean value was calculated and reported.

### Injury parameters

Information including the injury model (contusion, transection, etc.) and timepoint of sample collections was collected.

### Differential miRNA expression

Information including the miRNA extraction method, library generation, and detection platform was collected. Changes in miRNA levels was recorded as up- or down-regulated following SCI.

### Data analysis and synthesis

An initial synthesis of raw data was conducted to describe the studies in terms of initial results and methodological parameters of the primary literature. A meta-analysis was performed for all outcome measures. We applied the inverse variance meta-analysis using R package “meta” to aggregate available effect sizes (i.e. fold change) and standard deviation of the effect sizes for each miRNA. For studies where the authors only reported effect size and p-value, we extrapolated the standard deviation using the p-values and the inverse normal transformation. For miRNAs that only show up in one study, we simply report the effect size, extrapolated standard deviation and p-values; for miRNAs that show up in multiple studies, we reported the results based on the meta analysis. Finally, Benjamini-Hochberg Procedure was performed to calculate adjusted p-values. To examine the biological functions of the miRNAs, we performed functional enrichment analyses using miRNet 2.0 KEGG, REACTOME, and Gene Ontology (GO) pathway databases to identify miRNA targets of at least two miRNAs significant targets and pathways were considered for an adjusted p-value <0.05.

### Risk of bias

The use of SYRCLE’s risk of bias tool and the CAMARADES checklist for study quality was included to assess risk of bias and study quality by two reviewers. Any disagreement was resolved by consensus.

## Results

It would be valuable for identified SCI biomarkers to exist across species to facilitate the translation of therapeutic targets for SCI from bench to bedside. The growing body of literature investigating miRNAs as specific indicators of SCI pathophysiology represents a promising resource for the identification of intra-species biomarkers, that may have powerful representation of the tissue, time course, and mechanism of injury. We posited that an integrated understanding of SCI-specific miRNA changes could be achieved by synthesizing the existing literature produced over the last two decades. An overview of our experimental design is shown in Figure 1.

**Figure 1.**
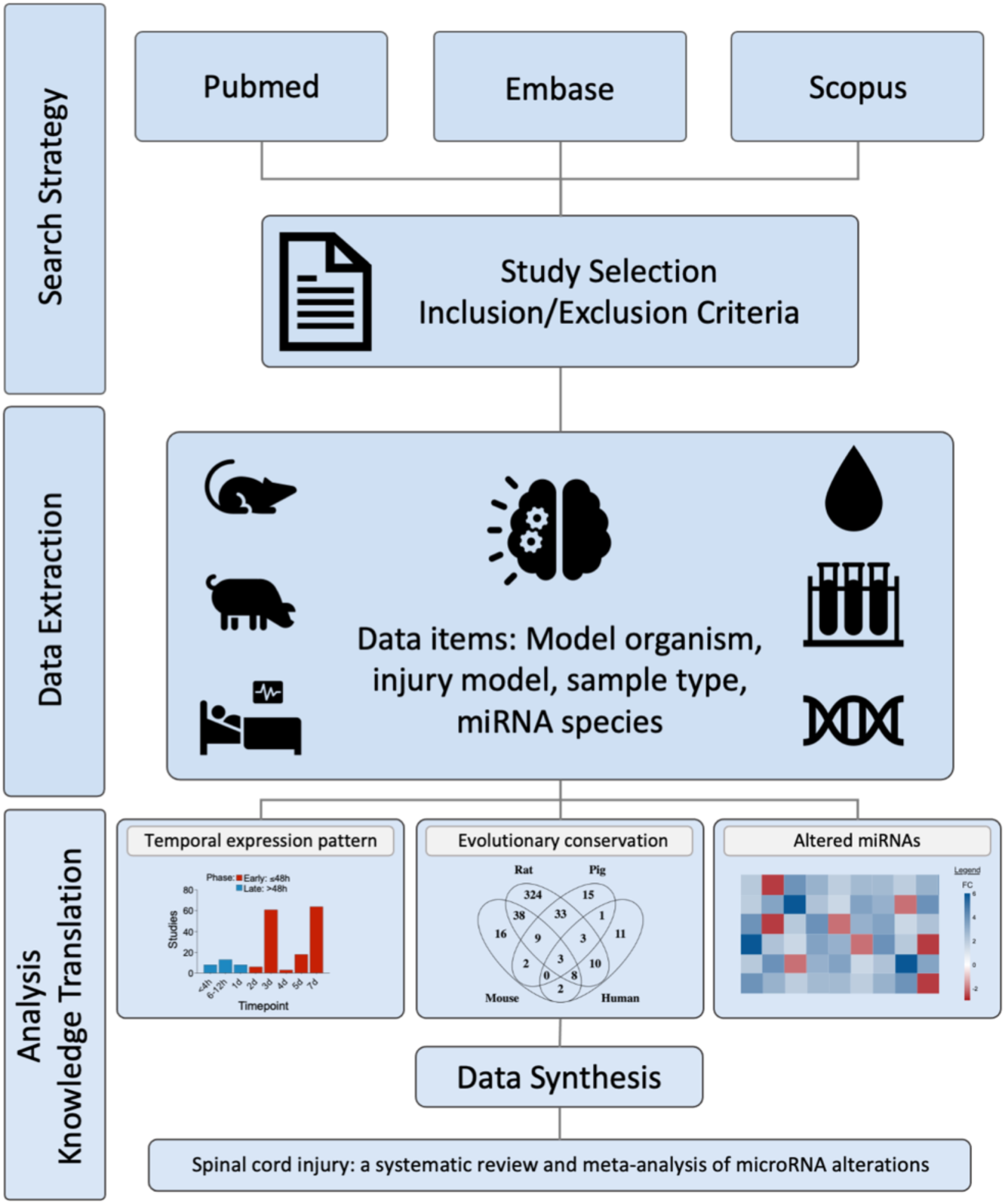
Study outline. Schematic overview of systematic approach to integration of studies of pathophysiology of microRNA changes following SCI.

### Study characteristics

We conducted a systematic analysis of the SCI literature describing miRNA changes and identified a total of 1878 studies eligible for screening from PubMed, Embase, and Scopus. A total of 1878 studies were identified for screening (Figure 2). Among these records, 939 were duplicate studies and were excluded. A total of 937 study titles and abstracts were screened, and 601 were excluded due to not meeting inclusion/exclusion criteria. Full-text review of the remaining 336 studies resulted in the exclusion of 202 studies. A total of 134 studies met inclusion/exclusion criteria and were included for analysis. The characteristics of studies included in this analysis are presented in Supplemental Table 1.

**Figure 2.**
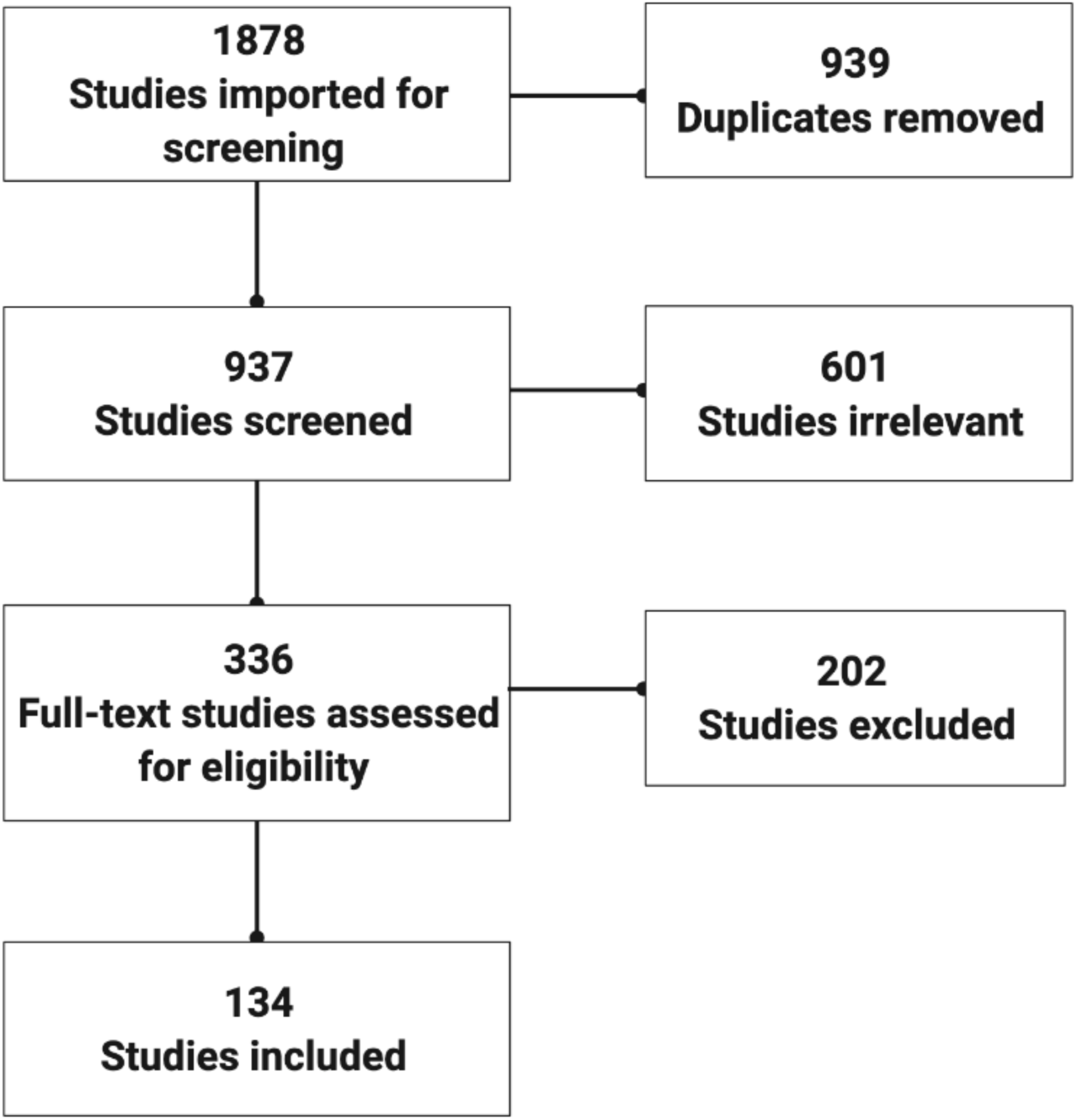
Flow diagram of study selection for meta-analysis.

We sought to characterize the current status of the body of knowledge of miRNA changes post-SCI with respect to model organism, injury model, molecular detection techniques and post-SCI time points studied. In addition to human patients (*H. sapiens*; six studies) with acute traumatic SCI, there were five species of animal models of SCI, including axolotl (*A. mexicanum*; two studies), Mouse (*M. musculus*; 32 studies), Rat (*R. norvegicus*; 80 studies, including Wistar and Sprague Dawley hybrids), and Pig (*S. scrofa*; four studies) (Figure 3A). SCI models included contusion (88 studies), compression (11 studies), transection (nine studies), and traumatic SCI (six studies) (Figure 3B). The molecular methods used to detect miRNA alterations included RT-PCR (105 studies), microarray (11 studies), and next-generation sequencing (NGS; six studies) (Figure 3C). Studies included in this systemic review detected a range of altered miRNA post-SCI, with some detecting a single altered miRNA, and others detecting hundreds of altered miRNAs. There were 47 studies that identified a single altered miRNA, 15 studies that identified two altered miRNAs, 14 studies that identified three altered miRNAs, 15 studies that identified four altered miRNAs, and 25 studies that identified five or more altered miRNAs (Figure 3D). Post-SCI changes were analyzed at time points ranging from one-hour post-injury to 90 days post-injury, with the majority focusing on three to seven days post-injury (Figure 4A).

**Figure 3.**
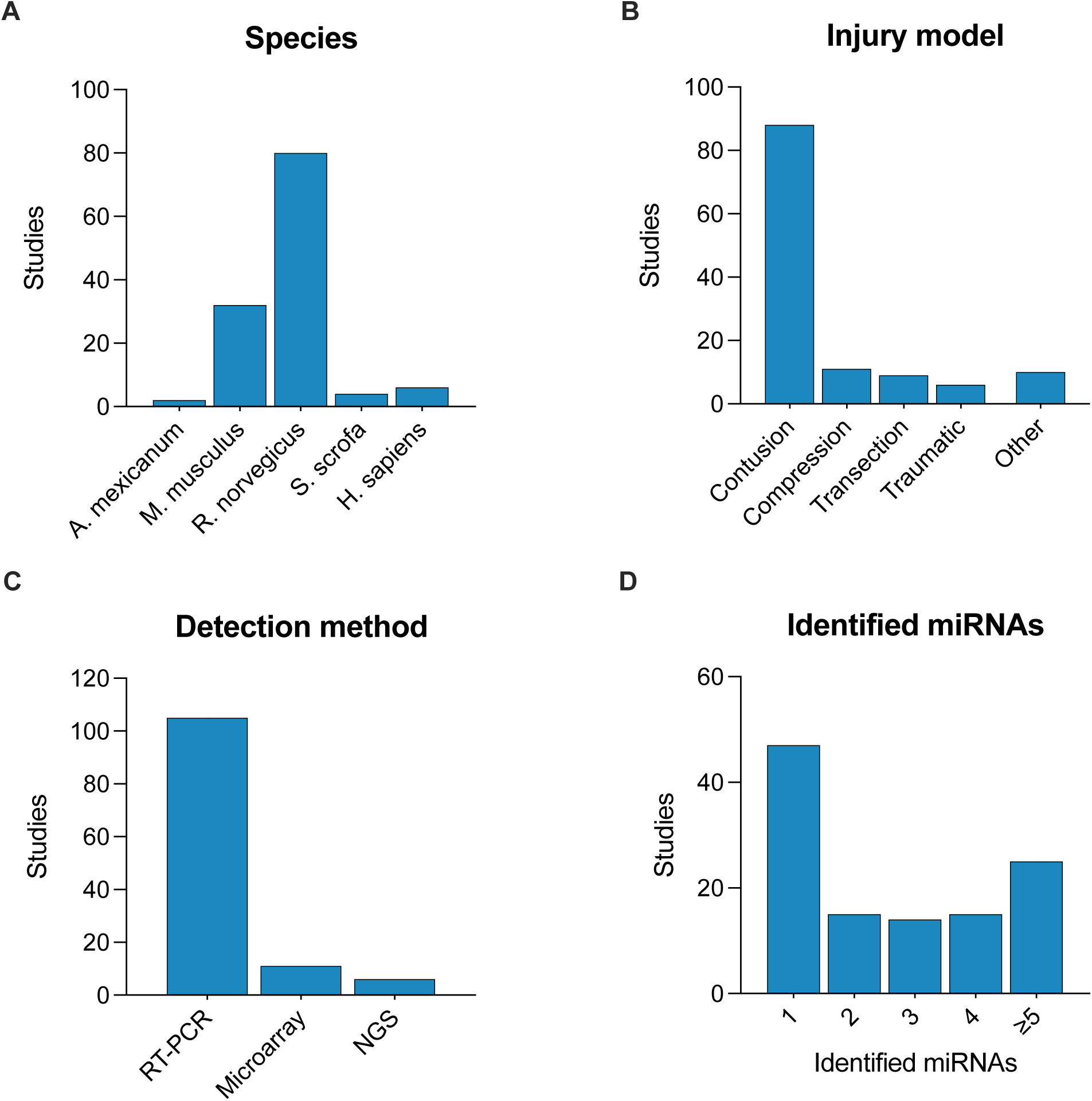
Characteristics of studies included in meta-analysis. Frequency of studies by A. Model organism. B. SCI injury model. C. miRNA detection method. D. Number of altered miRNAs detected. NGS: Next-generation Sequencing; miRNAs: microRNAs.

**Figure 4.**
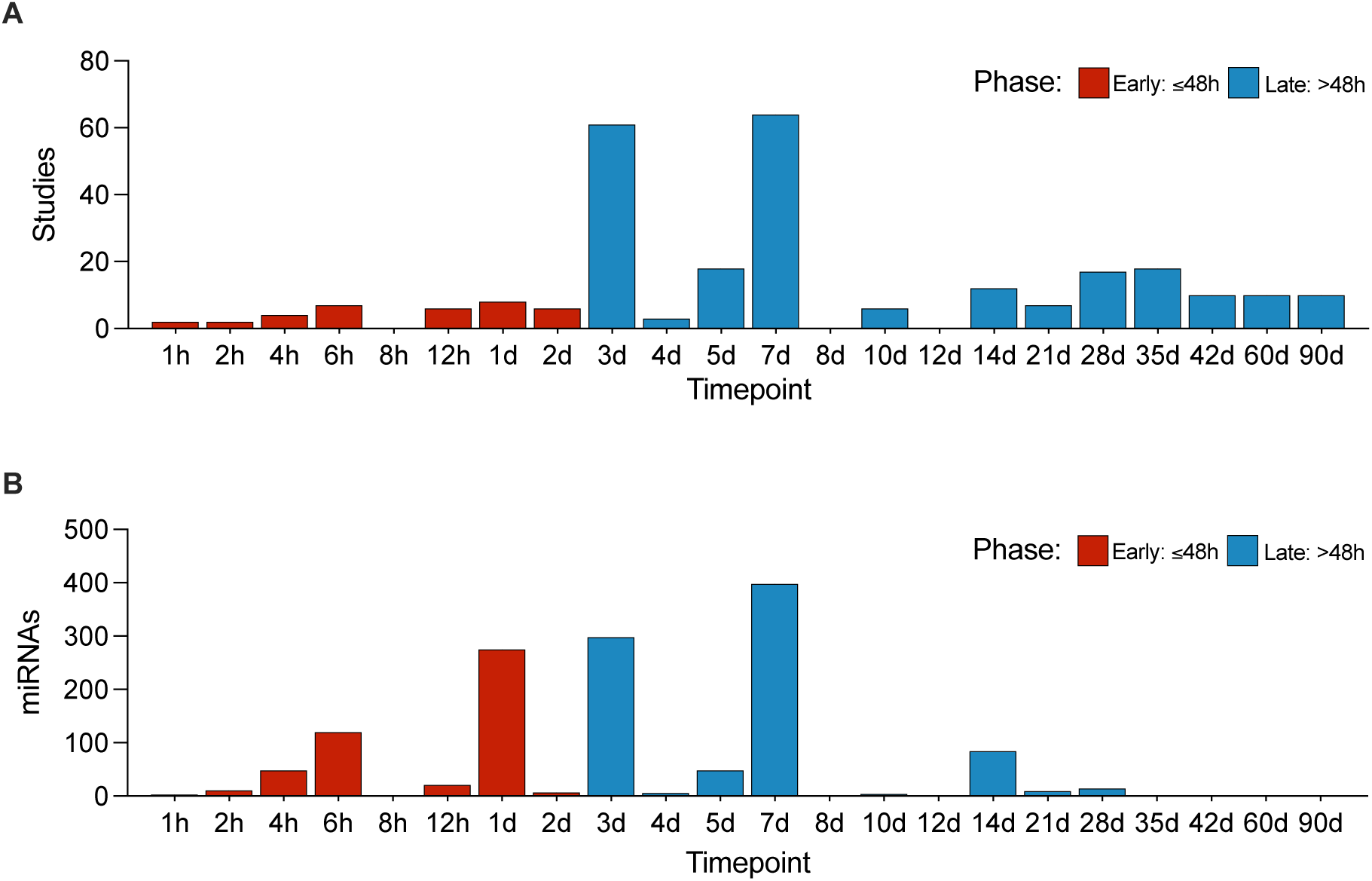
MicroRNAs implicated in the physiological response to SCI. Temporal characterization of A. study designs included in meta-analysis, and B. miRNA alterations.

### Systematic literature identification of miRNAs implicated in the physiological response to SCI

We sought to systematically establish the complete set of miRNAs identified during the physiological response to SCI. In total, 475 unique miRNAs were identified as altered during the pathophysiologic response to SCI across all studies. The number of miRNAs detected among the variety of study designs is shown in Supplemental Figure 1. A large proportion of altered miRNAs were detected in rat models of SCI (Supplemental Figure 1A), by studies using a contusion model of SCI (Supplemental Figure 1B), and by studies using high throughput methods like microarray and NGS (Supplemental Figure 1C). Among all miRNAs, 227 were implicated in the pathophysiology of SCI by more than one study and 34 miRNAs were implicated by more than five studies (Supplemental Figure 1D).

We next sought to identify the temporal pattern of altered miRNAs following SCI. miRNA alterations were assessed between one-hour and 90 days post-injury. The majority of altered miRNAs were identified within the first seven days post-injury, with the number of altered miRNAs highly related to the number of studies at a given time point (Figure 4A, B). Interestingly, the 24-hour timepoint was represented by only 3% of all studies, but the number of altered miRNAs at 24 hours represented 20% of all altered miRNAs across time points. Thus 24 hours post-injury may represent a biologically informative and potentially under-investigated time point for understanding the molecular response to SCI.

A critical knowledge gap in the literature is the identification of an integrated miRNA response to SCI *across* species, such that miRNA biomarkers and targets can be evaluated in preclinical animal models that can then inform clinical trials involving human patients. We sought to identify those miRNAs that are involved in SCI across multiple species (Figure 5). In total, 109 miRNAs were implicated in two or more species, 23 miRNAs in three or more species, and three miRNAs in four species, the list of which is provided in Supplemental Table 2.

**Figure 5.**
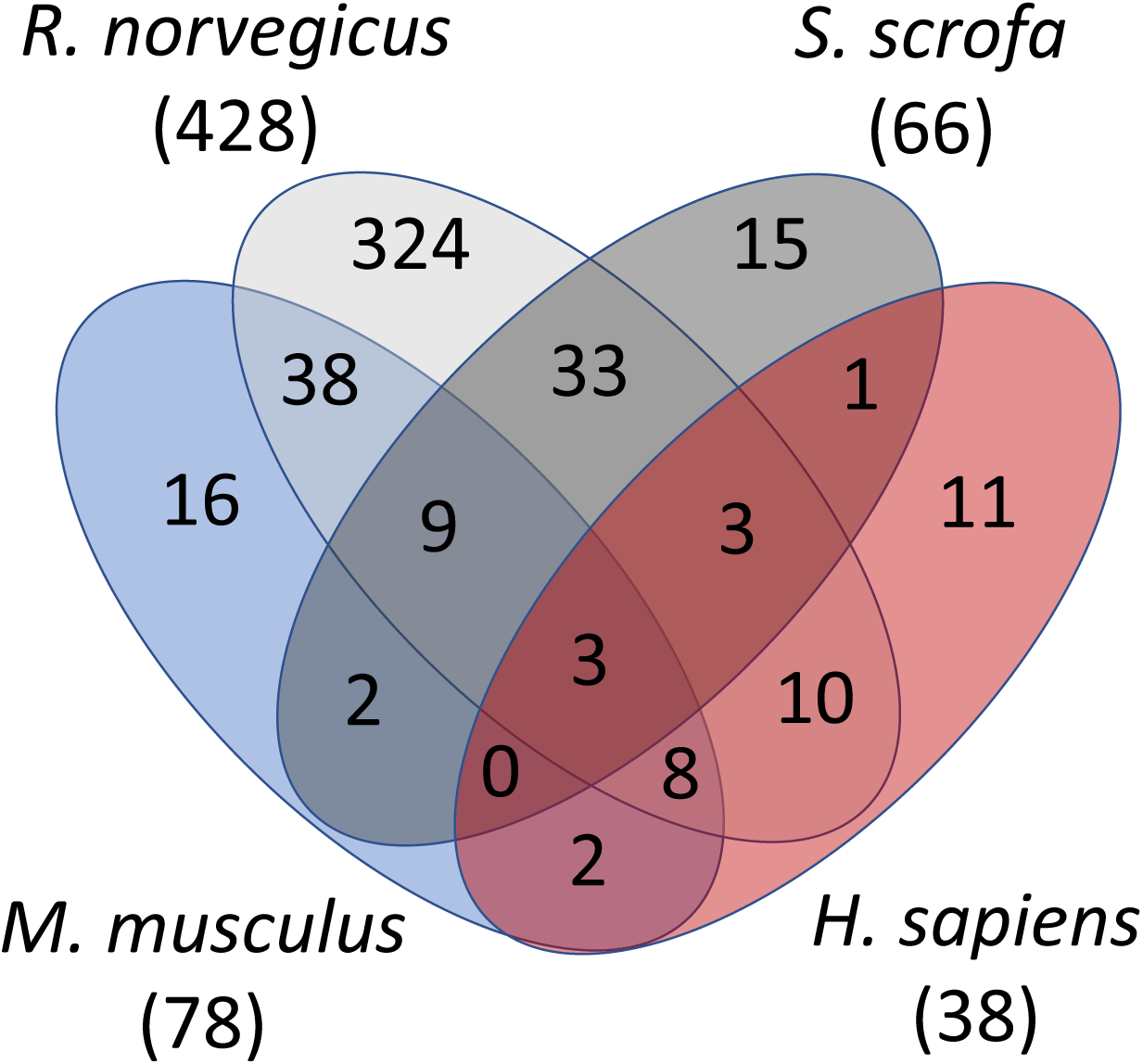
MicroRNA alterations across model organisms reveal an evolutionarily conserved miRNA response to SCI. Venn diagram showing altered miRNAs common and unique to each species. *M. musculus*: *Mus musculus*, mouse, blue; *R. norvegicus*: *Rattus norvegicus*, rat, grey; *S. scrofa*: *Sus scrofa*, pig; charcoal; *H. sapiens*: *Homo sapiens*, human, red.

### Meta-analysis reveals a tissue-targeted and temporally regulated miRNA response to SCI

We next sought to statistically integrate each small-scale experimental report of miRNA alterations post-SCI using an inverse variance meta-analysis using R package “meta” to aggregate available effect sizes (i.e. fold change) and standard deviation of the effect sizes for each miRNA. We first performed the analysis agnostic of species, tissue type, or time point, identifying a total of 98 altered miRNAs, of which 71 were upregulated (Table 1), and 27 were downregulated (Table 2). Next, we performed a subgroup analysis, seeking to identify temporally altered and/or tissue-targeted miRNAs that were identified by two or more studies. Time points were binned into two categories, “early” (≤48h) and “late” (>48h). First, analysis agnostic to tissue type revealed two miRNAs that were altered in the early post-SCI period, miR-21-5p and miR-223-3p, and 16 miRNAs that were altered in the late post-SCI period (Figure 6). The analysis of tissue-targeted miRNAs identified three miRNAs altered early following SCI in spinal cord parenchyma, and seven miRNAs altered in the late period. There were no miRNAs altered in the blood early following SCI, but four miRNAs were identified in the late period.

**Figure 6.**
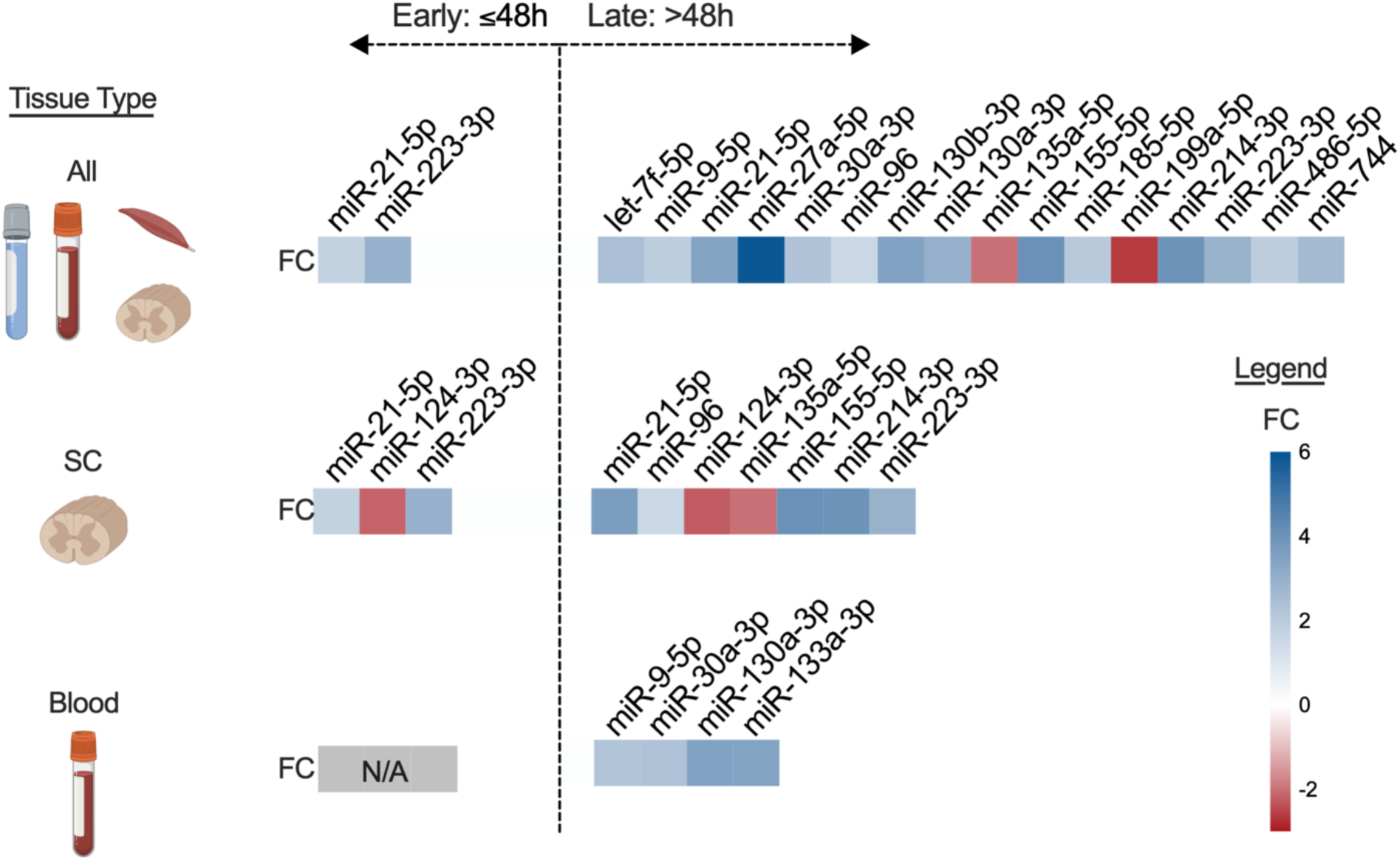
Meta-analysis of microRNAs implicated in SCI pathophysiology by two or more studies reveals a robust tissue-targeted and temporal miRNA response to SCI. SC: Spinal cord; FC: Fold Change.

**Table 1.**
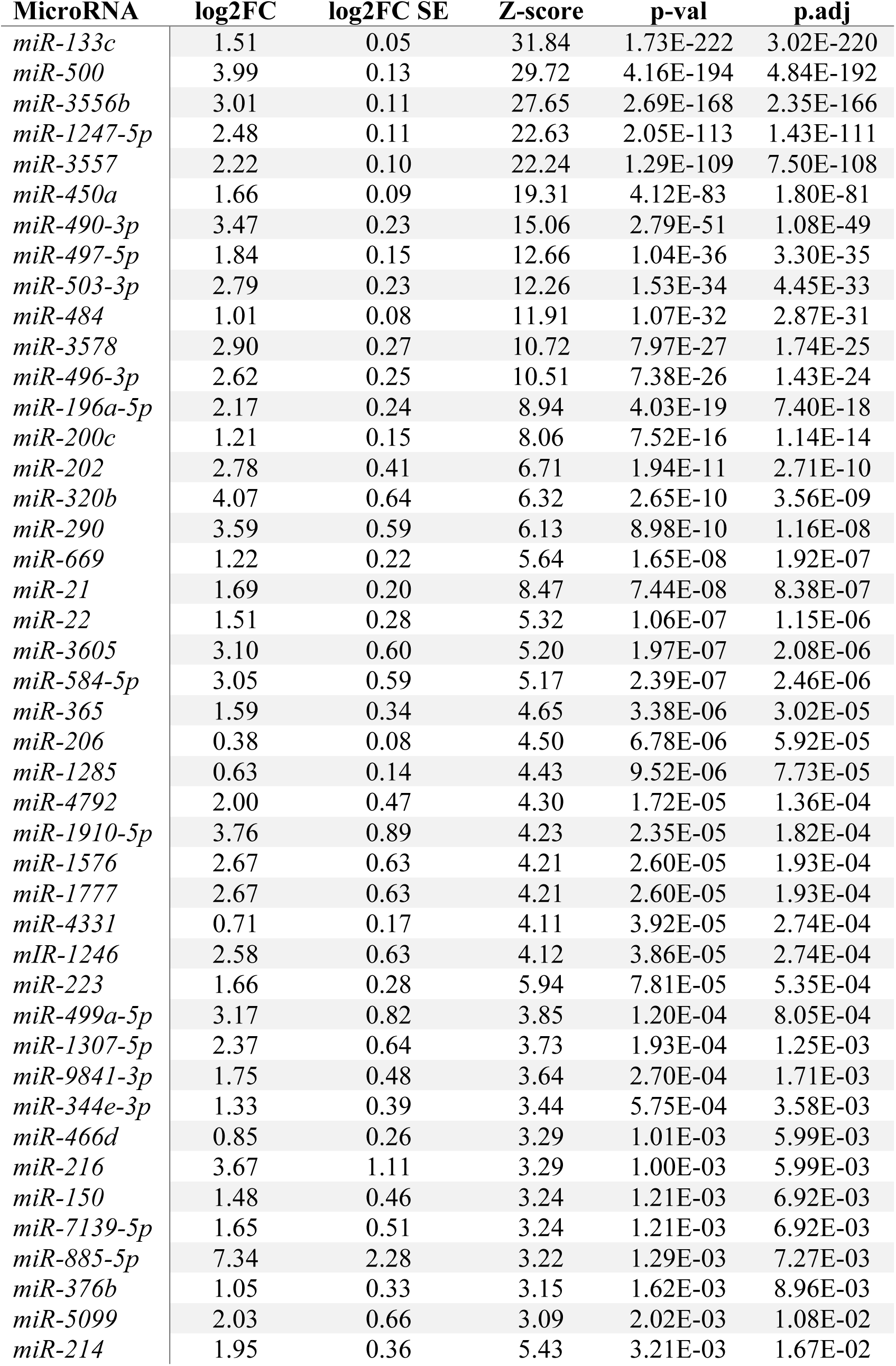
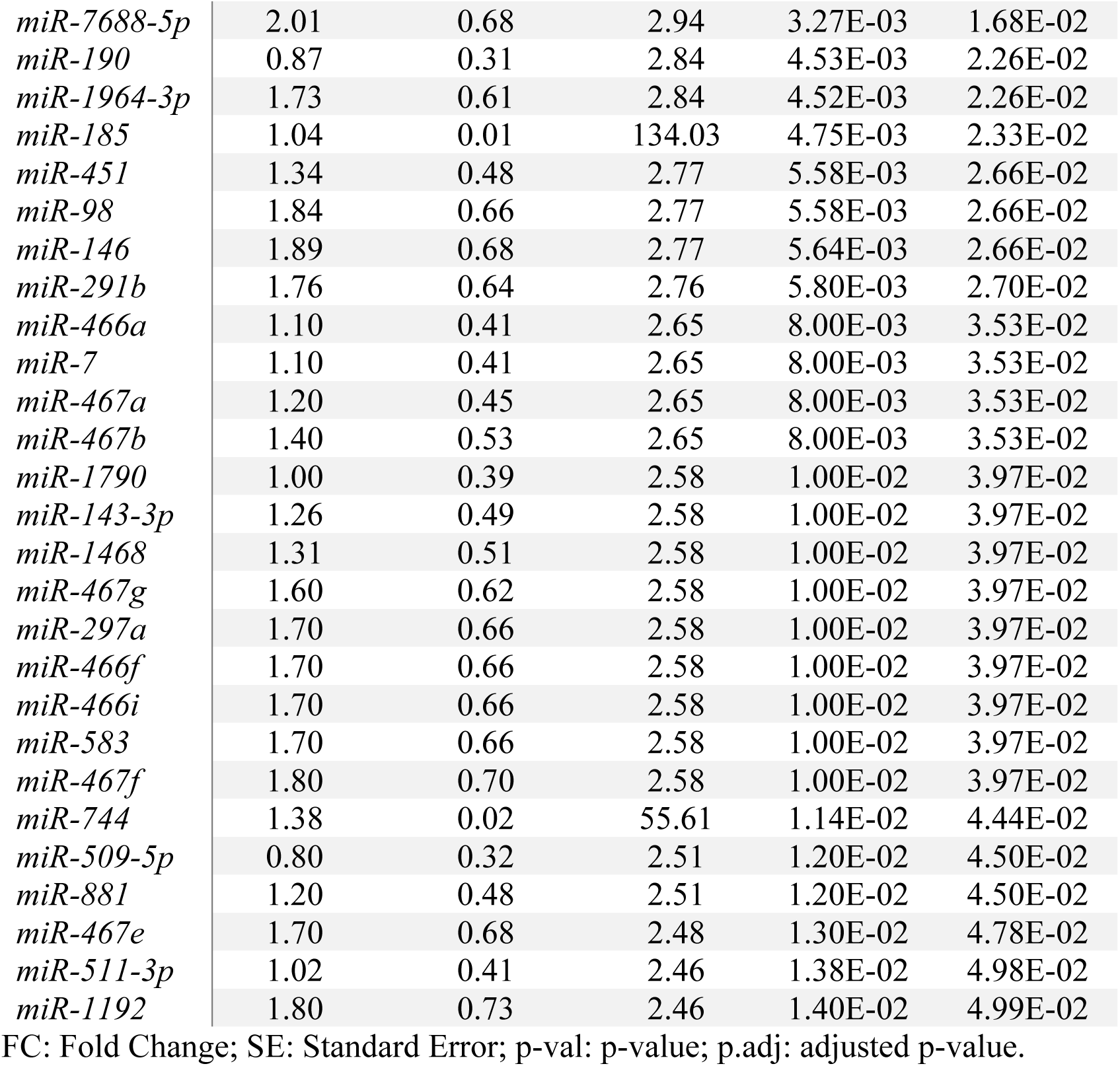
MicroRNAs upregulated following SCI.

**Table 2.**
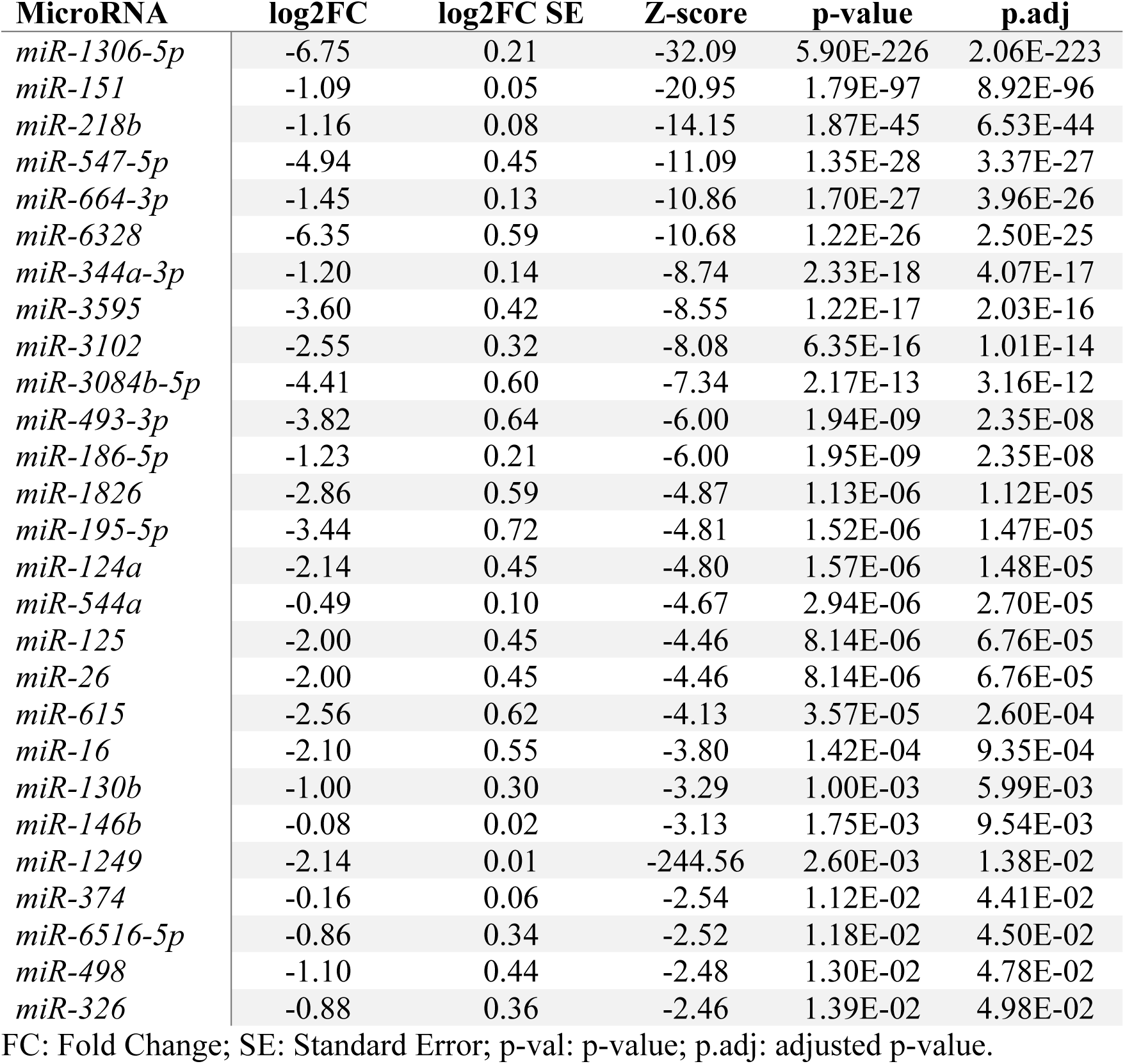
MicroRNAs downregulated following SCI.

### Functional characterization of altered miRNAs

To begin to characterize the biological significance of the altered miRNAs identified by meta-analysis and the roles they may play in the response to SCI, an interaction network was constructed between altered miRNAs and genes that were targeted by at least two miRNAs (Figure 7). KEGG and REACTOME pathway enrichment analysis revealed several significant pathways that are implicated in SCI processes including cellular stress response, cell death, axon guidance, and signaling molecules such as neurotrophin, TGF-beta, WNT and VEGF (Supplementary Tables 3, 4). Gene Ontology (GO) analysis, including biological process (BP), molecular functions (MF), and cellular compartments (CC) were determined, highlighting many systems relevant to the post-injury processes of SCI (Supplementary Tables 5, 6, 7).

**Figure 7.**
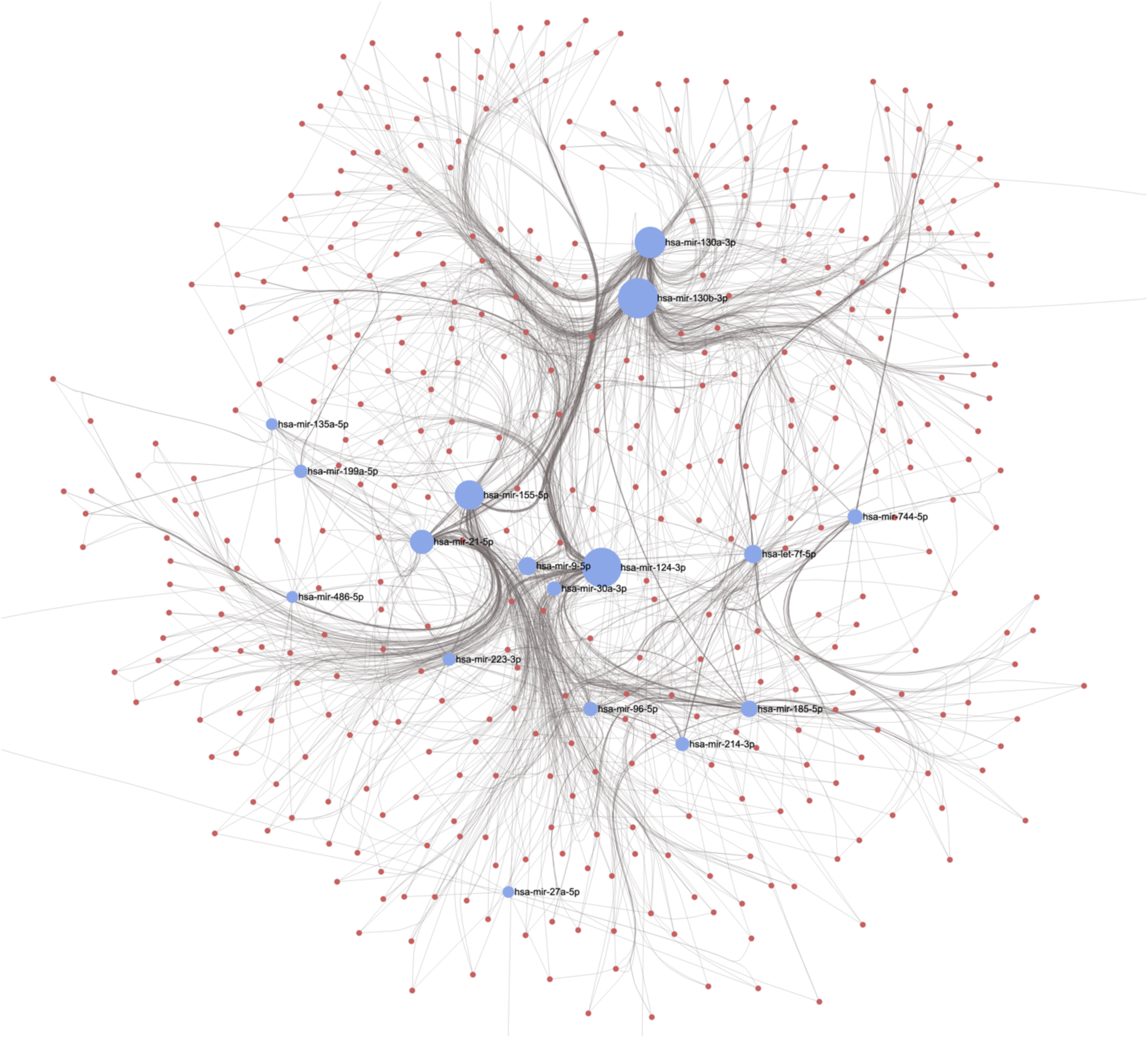
Network analysis of microRNAs identified by meta-analysis. Blue circles represent altered miRNAs. Red circles represent miRNA targets. The network was generated using miRNet web tool to explore altered miRNAs and their relationships and collective functions.

## Discussion

The mechanisms involved in the pathogenesis of SCI include a primary mechanical injury (impact) and a secondary injury induced by multiple subsequent biological processes, including a local inflammatory response, cytotoxicity, apoptosis, and demyelination [21, 40]. In addition to local inflammation, a systemic inflammatory response inducing organ damage has been shown to occur following SCI [32]. Although altered gene expression significantly contributes to the pathogenesis of secondary SCI [40], the regulatory networks that control it are not well understood. One aspect of the complex nature of secondary SCI could relate to gene regulation by miRNAs [20, 24, 25, 30]. As potential biomarkers of a pathological state, miRNAs are not constrained by cell membranes and communicate in extracellular fluids as free-floating miRNA [27] or within exosomes [26], and are considered stable, with relatively long half-lives of greater than 24 hours [19]. miRNAs are therefore powerful candidates for monitoring CNS pathophysiology related to SCI. In this study, we integrated and synthesized two decades of literature investigating miRNA changes following SCI. By integrating multiple miRNA expression datasets, from both experimental models and human patients with SCI, we identified a robust temporal pattern of miRNA alterations that appear to be tissue-targeted and conserved across multiple species. Our results provide a framework to understand the diverse, coordinated processes of miRNA regulation, within and beyond the spinal cord, following SCI.

A major challenge to the translation of preclinical therapies for acute SCI from bench to bedside is the relative lack of standardized physiologic assessments to enroll and stratify patients in large clinical trials [34]. Additionally, there are no established evolutionarily conserved molecules with which to benchmark therapeutic efficacy in preclinical animal models of SCI. Objective biomarkers for SCI that are conserved between humans and preclinical models including mouse, rat, and/or pig would be invaluable to the pursuit of SCI therapies [2, 4, 5, 11, 12, 15, 17]. In this study we identified miRNAs regulated following SCI that were conserved across species. These findings have several implications for the discovery and translation of new SCI therapies. The identification of animal models and associated surrogate markers of SCI development can facilitate rapid assessment of therapeutic efficacy before embarking on clinical trials. Without reliance on behavioral outcome measures that may not relate to the human condition, biomarkers that are consistently altered across species not only could reduce the number of study subjects, but could also improve confidence that preclinically discovered therapies might induce a promising response in clinical trials.

While highly effective therapies are currently lacking for patients with SCI, miRNAs represent promising targets to mitigate pathophysiologic process following injury [13]. In the meta-analysis presented here, we provide a comprehensive resource to the SCI community both by quantifying where current evidence is strong, and by identifying where there are gaps in knowledge. For example, there is a paucity of data describing miRNA changes in human SCI patients, and limited studies providing a comprehensive analysis of the time course of miRNA changes following SCI. The identification of these gaps may serve as guidance for future investigations of SCI-related miRNAs. We also provide a tool for the identification of potential pathways to target in the development of novel therapies. Finally, we synthesize the miRNA changes in over 130 studies, identified those that were conserved across species, timepoints, and tissue types, and report their predicted targets and pathway involvement.

This is a critical juncture in the nascent field of miRNA biology in SCI, at which a synthesis and rigorous objective analysis of the collective data amassed with respect to miRNA changes following SCI can significantly inform future investigations. This systematic review and meta-analysis is intended to serve as a resource to the SCI community, providing a comprehensive clearinghouse of robust miRNA changes across a spectrum of SCI paradigms with various injury models, species, tissues, and time points. This systematic review and meta-analysis establishes a thorough, rigorous, and unbiased window into miRNAs and their targets to enable the discovery of the next generation of biomarkers and therapeutic interventions.

### Strengths and limitations

Here we present a comprehensive and systematic analysis of miRNA changes following SCI that seeks to assist researchers and clinicians in the pursuit of biomarkers and novel therapeutics for SCI. The main limitations of this study include the heterogenous nature inherent to the diversity of studies included in the meta-analysis presented here. A wide range of model organisms, injury models, time points, and detection techniques are included. Additional potential limitations include the exclusion of possible articles published following our analysis but prior to its publication, not indexed in PubMed, Embase, or Scopus, and of those articles that might not include relevant keywords or phrases used in this search.

### Conclusions

This systematic review and meta-analysis provides a comprehensive description of the changes that occur across miRNAs during the early and late post-injury phase of acute SCI. Here, we identify miRNAs that are tissue-restricted and evolutionarily conserved across species, providing a resource of robust miRNA alterations that play biological roles in the pathophysiologic processes of SCI. These miRNAs represent promising candidates for biomarkers of SCI diagnosis and prognosis, as well as potential targets for future SCI therapies. With the emergence of future SCI therapies seeking validation in clinical trials, the field of SCI is in need of additional approaches to classifying injury severity and improved methods of predicting outcome. miRNAs represent promising solutions to both a need for biomarkers and the bottleneck in the pipeline from the bench to developing clinically relevant therapies for SCI.

## List of abbreviations

CNS: Central nervous system
CSF: Cerebrospinal fluid
miRNA: microRNA
SCI: Spinal cord injury

