## Supplementary figures and images for "Spinal cord injury: a systematic review and meta-analysis of microRNA alterations"

### Supplemental Figures

**Figures and Tables**

Figure 1. Impact of study design on number of altered miRNAs detected

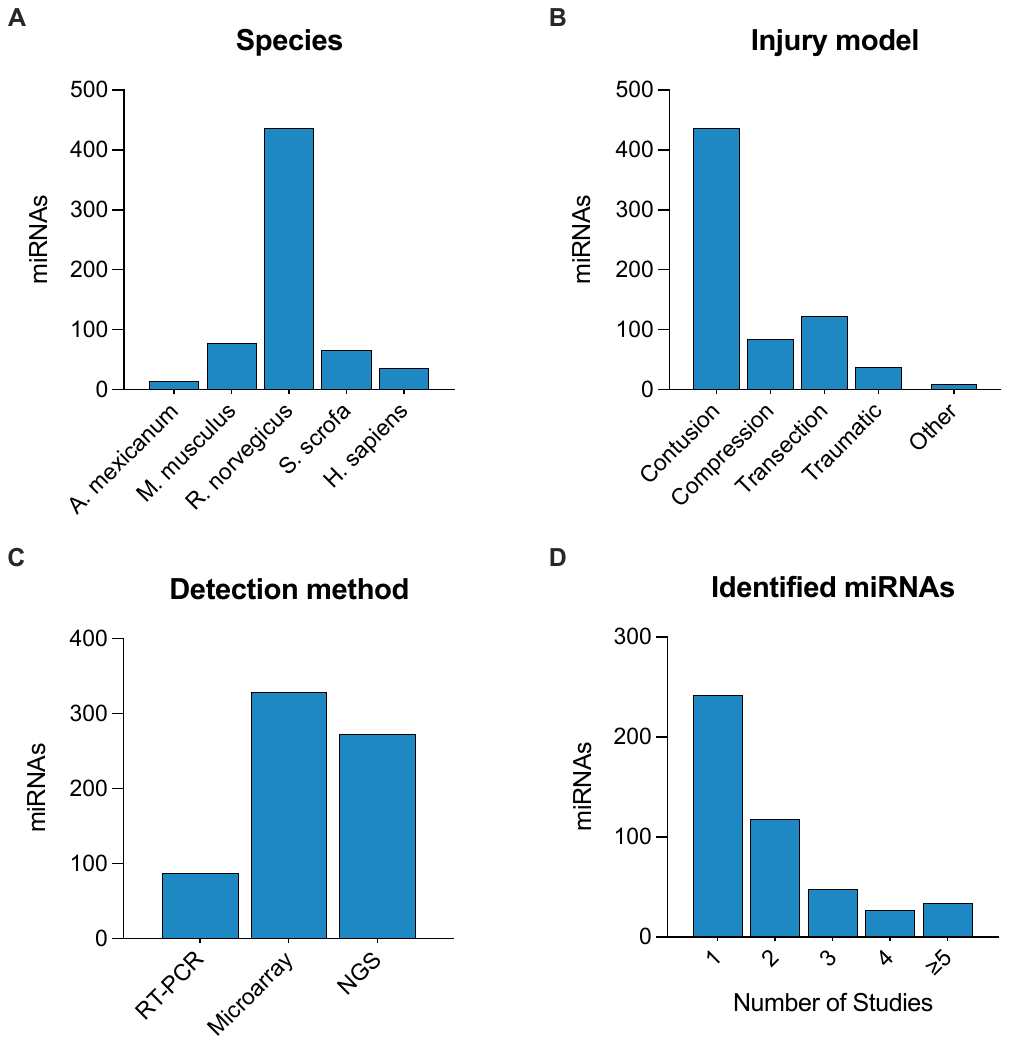
